# Role of the default mode network in cognitive transitions

**DOI:** 10.1101/295683

**Authors:** Verity Smith, Daniel J Mitchell, John Duncan

## Abstract

A frequently repeated finding is that the default mode network (DMN) shows activation decreases during externally-focused tasks. This finding has led to an emphasis in DMN research on internally-focused self-relevant thought processes. A recent study, in contrast, implicates the DMN in substantial externally-focused task switches. Using functional magnetic resonance imaging, we scanned 24 participants performing a task switch experiment. Whilst replicating previous DMN task switch effects, we also found large DMN increases for brief rests as well as task restarts after rest. Our findings are difficult to explain using theories strictly linked to internal or self-directed cognition. In line with principal results from the literature, we suggest that the DMN encodes scene, episode or context, by integrating spatial, self-referential and temporal information. Context representations are strong at rest, but re-reference to context also occurs at major cognitive transitions.

---

The default mode network (DMN) is one of the most robust discoveries of neuroimaging. Default mode regions – prominently including parts of medial frontal cortex, posterior cingulate, posterolateral parietal cortex, retrosplenial cortex and hippocampal formation – show strong functional connectivity at rest (Greicius et al., 2009; Fransson & Marrelec, 2008; Andrews-Hanna et al., 2010) and commonly increase or decrease activity together across a wide range of cognitive manipulations (Shulman et al., 1997; Spreng et al., 2009). Despite these robust findings, the functional significance of the DMN remains unclear.

Following the early work of Shulman et al. (1997), one much-replicated result is decreased DMN activity in many tasks compared to rest. These decreases complement common patterns of task-related increase in other networks, including “dorsal attention” (Corbetta & Shulman, 2002) and “multiple-demand” or MD networks (Duncan & Owen, 2000; Duncan, 2010). Stronger activity during rest compared to focused task performance led early on to the proposal that the DMN is involved in internally-generated cognition, including mind-wandering (Mason et al., 2007; Christoff et al., 2009) and self-related thought (Johnson et al., 2002; D’Argembeau et al., 2005; Andrews-Hanna et al., 2010). Subsequent findings lend support to this emphasis on internally-directed and self-relevant cognition, including DMN activation during autobiographical memory recollection (Diana et al., 2007; Schacter et al., 2007; Vilberg & Rugg, 2012), imagining possible future events (Addis et al., 2007; Spreng et al., 2010; Andrews-Hanna et al., 2010), making self-referential judgments (Johnson et al., 2002; D’Argembeau et al., 2005), and imagining routes (Kumaran & Maguire, 2005; Spiers & Maguire, 2007; Howard et al., 2014; Balaguer et al., 2016) etc. A broad suggestion is that the DMN creates internal scenes (Hassabis & Maguire., 2007, 2009), episodes (Buckner & Carroll., 2007; Addis et al., 2007) or contexts (Bar, 2007, 2009; Ranganath & Ritchey., 2012), allowing cognition to escape from the constraints of the present environment. Such an internal scene or context might include spatial, temporal, social and perhaps other elements. Both imaging and animal experiments, for example, link parts of the DMN (especially posterior cingulate, retrosplenial cortex, hippocampus) to spatial representation and navigation (Vann & Aggleton, 2002; Burwell et al., 2004; Bachevalier & Nemanic, 2008; Doeller et al., 2010; Jacobs et al., 2013; Spiers & Maguire, 2007; Howard et al., 2014; see Bird & Burgess., 2008 for a review). Medial frontal and posterior parietal regions show strong activity linked to social cognition, including consideration of others’ mental states (Frith & Frith, 2003; Kumaran & Maguire, 2005; Amodio & Frith, 2006). Much DMN activity is also linked to time, as in recollection and future planning (Addis et al., 2007; Andrews-Hanna et al., 2010).

Recently, however, a number of results raise questions over a distinction between processes involved in external and internal cognition (Lee et al., 2005; Christoff et al., 2009; Baldassano et al., 2016; Chen et al., 2017). Of most relevance to the current work, a study by Crittenden et al. (2015) implicated the DMN in externally-focused task switching. In this study, participants were asked to perform a yes/no task following a task rule cued for by the color of frame surrounding the imperative stimuli. Two tasks were associated with each of three stimulus domains (pictures, words and shapes). Tasks were presented sequentially in a pseudorandom order to create stay trials (where the current task is the same as the previous task), within-domain switch trials (where the current task involves the same stimulus domain as the previous task) and between-domain switch trials (where the current task involves a different domain of stimulus compared to the previous task). Following Andrew-Hanna et al. (2010), Crittenden et al. (2015) divided the DMN into 3 subnetworks, “core” (anteromedial frontal cortex and posterior cingulate), “medial temporal (MTL)” (retrosplenial, parahappicompal and hippocampal cortex, along with posterior inferior parietal and ventromedial frontal cortex) and “dorsomedial prefrontal (dmPFC)” (dorsomedial prefrontal cortex accompanied by regions in lateral temporal lobe and temporo-parietal junction). Contrary to the common finding of decreased DMN activity during demanding, externally-focused cognition, Crittenden et al. (2015) found core and MTL subnetworks to increase activity during the most demanding, between-domain switch trials (with the dmPFC subnetwork, if anything, showing the reverse). In all 3 subnetworks, furthermore, multivoxel pattern analysis (MVPA) showed distinct activity patterns for different trial types, in particular for the three different stimulus domains. Apparently, DMN activity can be seen not only in internally-directed cognition, but in some aspects of external task switching.

Here we followed up this lead to DMN function. In a study similar to that of Crittenden et al. (2015), we asked participants to switch between trials involving multiple rules and stimulus domains. There were two tasks associated with each of the 3 stimulus domains creating three types of task switch: between-domain switch, within domain switch and stay trials. To relate task switching activity to standard “rest” activity, we included cued rest trials in which participants simply rested, waiting for the next trial to begin. Rest trials could follow other rest trials, creating ‘rest stay’ trials, or follow task trials creating ‘rest switch’ trials. Importantly, these rests were short, of the order of the duration of a single trial, giving participants little time for explicit mind-wandering. Incorporation of rest trials, furthermore, allowed us to examine the switch back from rest to task (restart trials). To separate task preparation from execution, we introduced a delay between the colored cue instructing which task to perform next, and the imperative stimulus allowing the task to be executed. The experiment design is presented in Figure 1 for a full description, see the Experimental Procedures. Following on from Crittenden et al. (2015), we assessed univariate switch and rest-related activity and task-related multivariate activity patterns in previously defined core, MTL and dmPFC subnetworks Andrews-Hanna et al. (2010), as well as the typical task-positive MD network (Fedorenko et al., 2013). Our results are inconsistent with views of DMN function strictly linked to internal or self-directed cognition. In line with many others, we propose that the DMN indeed encodes scene, episode or context, but assign this context encoding a direct role in implementation and control of current, externally-focused cognition as well as internal thought processes.

**Figure 1.**
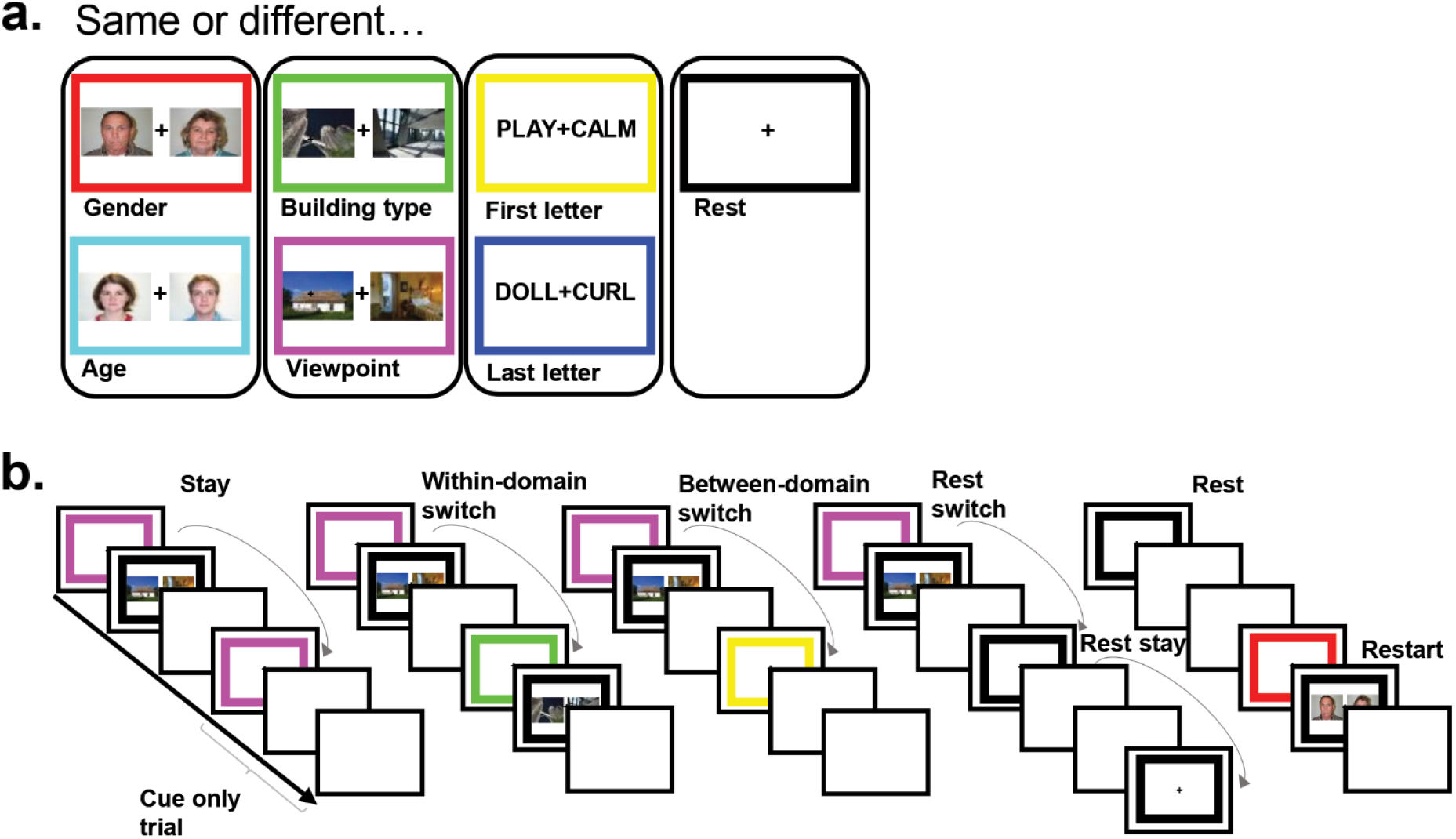
Task design. Participants were required to make same/different judgements on pairs of stimuli based on a task rule. Each task rule was cued by the frame color, learnt by participants in a training session prior to scanning. a. Each of the six tasks and their associated frame color. There were 3 stimulus domains with 2 task rules associated with each. An additional black frame cued rest trials in which there was no upcoming task to complete. b. Experimental design. Each trial consisted of a 2 second cue phase in which the colored frame specifying the task rule for the upcoming trial (or rest trial) was presented, followed by an execution phase (until response) or a 1.2 second delay, followed by a 1.75 second intertrial interval. ‘Cue-only trials’ refer to task trials where there was no execution phase. The tasks were presented in a pseudorandom order creating 6 switch conditions (stay trials (task trials preceded by the same task), within-domain switch (task trials preceded by same domain task trials), between-domain switch (task trials preceded by different domain task trials), rest switch (rest trials preceded by task trials), rest stay (rest trials preceded by rest trials) and restart (task trials preceded by rest trials).

## RESULTS

### Behavioral switch costs

Participants performed with an average of 95.9% correct responses (SD = 0.03). Paired-sample t-tests showed that responses were significantly faster for stay trials (1217ms) compared to within-domain switch trials (1373ms, t(23) = 4.87, p < 0.01), and between-domain switch trials (1348ms, t(23) = 3.97 p < 0.01), with no significant difference between the two types of switch trial.

### Increased DMN activity on rest trials

Given our interest in cognitive switching, our fMRI analyses focused largely on cue-related activity, with activity for task execution removed (see Experimental Procedures). Our first analysis tested for the typical “task-negative” characteristic of the DMN, with stronger activity during rest compared to task. To this end, for each DMN subnetwork, we compared cue activity on rest and task trials, the latter defined as the mean of stay, within-domain switch and between-domain switch trials. For all univariate analyses, average contrast values for each region of interest (ROI) were extracted using the MarsBaR SPM toolbox (Brett et al., 2002), and contrast values were then averaged across ROIs within each subnetwork (see Experimental Procedures). Figure 2 shows average contrast values for rest switch>task and rest stay>task. There were significant increases in activity during rest switch compared to task in all DMN subnetworks (core: t(23) = 3.45, p < 0.01; MTL: t(23) = 4.84, p < 0.01; dmPFC: t(23) = 2.34, p < 0.05), and for rest stay compared to task in the core and MTL subnetworks (core: t(23) = 3.34, p < 0.01; MTL: t(23) = 4.88, p < 0.01), but not in the dmPFC subnetwork (t(23) = 1.46, p > 0.05). Additional t-tests revealed increased activity in rest stay compared to rest switch trials only in core and MTL subnetworks (core: t(23) = 2.09, p < 0.05; MTL: t(23) = 3.16, p < 0.01; dmPFC: t(23) = 0.36, p > 0.05). We ran a two-way repeated measures ANOVA with factors contrast (rest stay>task stay, rest switch>task stay) and DMN subnetwork (core, MTL, dmPFC) to test for an interaction effect between rest switch type and DMN subnetwork. While the effect of contrast was not significant (F(1,23) = 3.88, p >0.05), significant main effects of subnetwork (F(2,46) = 12.53, p < 0.01) and a significant interaction effect were found (F(2,46) = 8.72, p < 0.01).

**Figure 2.**
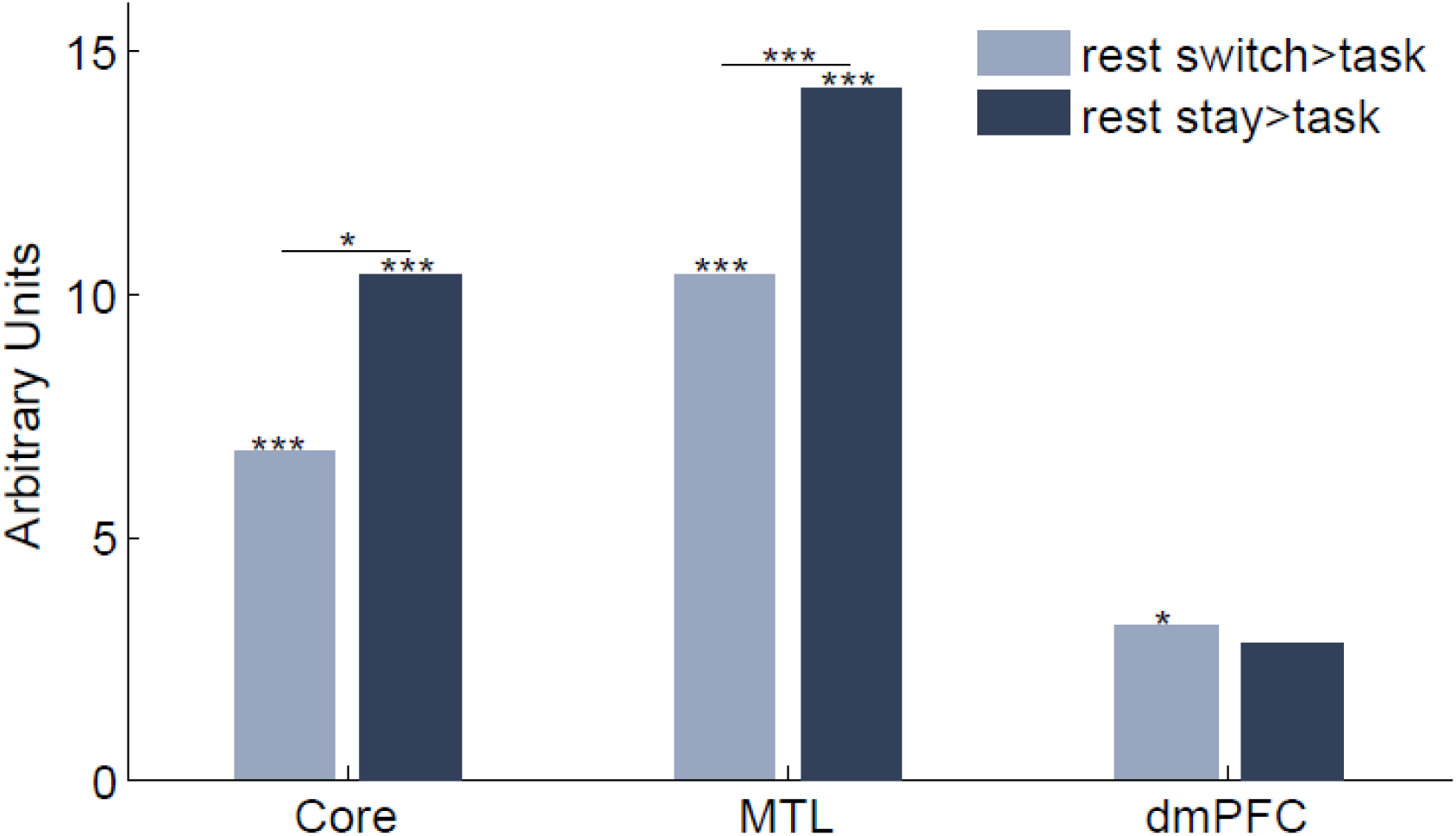
Contrasts of rest compared to task activity in each DMN subnetwork. Average contrast values for rest switch (light bars) and rest stay (dark bars) for each DMN subnetwork compared to mean of all task cue periods. Significant (p < 0.05) increases in rest activity compared to task, as well as paired t-tests between contrasts within DMN subnetworks, are indicated with * = p < 0.05, ** = p < 0.02, *** = p < 0.01.

### Increased DMN activity for large task switches

Second, we aimed to replicate the results of Crittenden et al. (2015), showing increased activity in core and MTL subnetworks for between-domain switch trials. Figures3a-c show the effects of switch condition on cue period activity in each DMN subnetwork. Core and MTL DMN subnetworks showed increased activity for between-domain switches compared to both within-domain switches (core: t(23) = 2.16, p < 0.05; MTL: t(23) = 2.25, p < 0.05) and stay trials (core: t(23) = 2.40, p < 0.05; MTL: t(23) = 2.46, p < 0.05). In line with trends reported by Crittenden et al. (2015), the dmPFC subnetwork showed the opposite effects of switch type, with decreased activity for between-domain switches compared to within-domain switches (t(23) = 2.14, p < 0.05). Within-domain switch trials were not significantly different from stay trials and in a supplementary analysis, no significant effects of switch type were found during the execution phase for core and MTL DMN subnetworks while the dmPFC subnetwork showed decreased activity for between-domain switch trials compared to within-domain switch (t(23) = 2.55, p < 0.02) and stay (t(23) = 3.13, p < 0.01).

**Figure 3.**
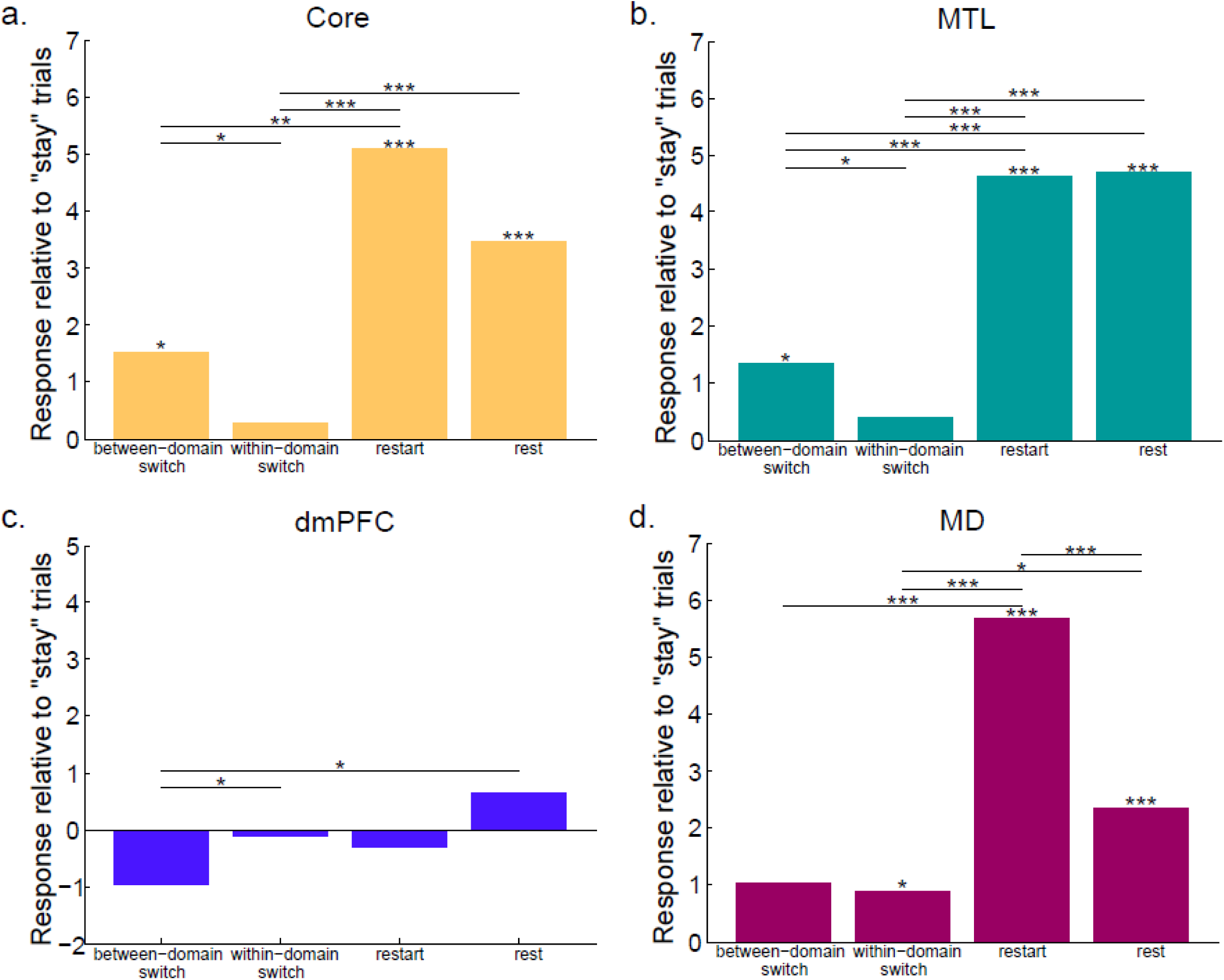
Contrasts of between-domain switch, within-domain switch, restart and rest compared to task stay trials for each DMN subnetwork and MD regions. (a. Core DMN, b. MTL DMN, c. dmPFC DMN, d. MD). Significant (p < 0.05) increases in activity compared to stay, as well as paired t-tests between contrasts within (sub)networks, are indicated with * = p < 0.05, ** = p < 0.02, *** = p < 0.01.

### Large increases in DMN activity for task restarts

Finally we were interested in restart activity, i.e. activity on trials where participants switched back from rest to task. For each DMN subnetwork, Figures 3a-c also show the contrast values for restart>stay, with rest>stay (mean of rest stay and rest switch trials) added for comparison. For core and MTL, but not dmPFC, subnetworks, t-tests revealed increased activity during both restart (core: t(23) = 3.87, p < 0.01; MTL: t(23) = 3.92, p < 0.01; dmPFC: t(23) = 0.38, p > 0.05) and rest (core: t(23) = 4.42, p < 0.01; MTL: t(23) = 5.38, p < 0.01; dmPFC: t(23) = 1.15, p > 0.05) compared to stay.

Our design allowed for separation of cue and execution-related activity in stay, within-domain switch and between-domain switch trials. This was achieved by including cued task trials without an execution phase, reducing the covariance between the two task phases. However, all restart trials were full trials including an execution phase. As such, our design could not separate cue and execution components in this condition. To check our conclusions in relation to restart-related activity, we repeated analyses using beta values from whole trials (defined as the average beta values of both cue and execution phases of restart and stay trials). These analyses similarly found increased activity for restart compared to stay activity in core and MTL DMN subnetworks (core: t(23) = 5.00, p < 0.01; MTL: t(23) = 5.95, p < 0.01) and increased activity during rest compared to stay in the core DMN subnetwork (t(23) = 7.63, p < 0.01). MTL rest activity was not greater than whole stay trial activity. In the dmPFC DMN subnetwork, rest activity was greater than stay (t(23) = 7.98, p < 0.01) but not restart.

### Similarities and differences between MD and DMN response profiles

To examine activity patterns in the MD network, we averaged results across canonical MD regions defined in lateral frontal, dorsomedial frontal, insular and parietal cortex (see Experimental Procedures). Figure 3d shows the average cue period contrast values for between-domain switch, within-domain switch, restart and rest conditions, all compared to stay trials. As in core and MTL subnetworks, t-tests showed greater MD activity for restart (t(23) = 4.34, p < 0.01) and rest trials (t(23) = 3.05, p < 0.01). MD activity was also greater in within-domain switch trials (t(23) = 2.20, p > 0.05) compared to stay trials. Although between-domain switch trials were not significantly different from stay trials, there was also no difference between within-domain switch and between-domain switches. Again, a supplementary analysis showed no significant effects of switch type during task execution.

To test whether differences in switch effect between subnetworks of the DMN and MD network were significant, we ran two two-way repeated measures ANOVAs with factors contrast (between-domain switch>stay, within-domain switch>stay) and network (core, MD) and (MTL, MD). The ANOVA comparing core DMN with MD activity found no significant main effects of contrast or network, but a significant interaction effect (F(1,23) = 5.21, p < 0.05) showing that the difference between between-domain switch and within-domain switch was greater in core DMN than MD regions. Likewise, the ANOVA comparing MTL DMN with MD activity also found no significant main effects of contrast or network, but a significant interaction effect (F(1,23) = 6.94, p < 0.02) showing that the difference between between-domain switch and within-domain switch was greater in MTL DMN than MD.

Another difference between MD and DMN networks is that, in MD regions, activity was significantly greater for restart trials – requiring active task performance – than for rest trials. To test this difference directly, we ran two two-way repeated measures ANOVAs with factors contrast (restart>stay, rest>stay) and network (core, MD) and (MTL, MD). The ANOVA comparing core DMN with MD activity found a significant main effect of contrast (F(1,23) = 10.36, p < 0.01) but not network, with a significant interaction effect (F(1,23) = 8.55, p < 0.01) showing that the difference between restart and rest was greater in MD regions than core DMN. The ANOVA comparing MTL DMN with MD activity also found a significant main effect of contrast (F(1,23) = 5.27, p < 0.05) but not network, with a significant interaction effect (F(1,23) = 29.35, p < 0.01) showing that the difference between restart and rest was greater in MD regions than MTL DMN.

### Individual voxels in DMN and MD regions show sensitivity to both rest and task restart

To test whether voxels within the same regions were sensitive to rest as well as to the large restart back to task, for each participant in each ROI we calculated the proportion of voxels showing above threshold responses to both rest>task and restart>stay contrasts, or either individually, at the threshold value of p = 0.05, uncorrected. The proportions of voxels sensitive to both contrasts and each individually were then averaged across participants and across (sub)network regions of DMN and MD cortex. All (sub)network regions showed voxels sensitive to both rest and restart (Core: 9.7%, MTL: 10.5%, dmPFC: 5.6%, MD: 10.1%) as well as voxels showing unique sensitivity to restart (Core: 9.2%, MTL: 8.6%, dmPFC: 6.2%, MD: 15.3%) and rest (Core: 8.6%, MTL: 12.5%, dmPFC: 9.0%, MD: 8.9%).

### DMN and MD activity patterns distinguish task domains

Multivariate analyses were also carried out to establish whether DMN cue period activity could distinguish between different task types. For each task pair (e.g. age vs. building type), a support vector machine (LIBSVM; Fan et al., 2005) was trained to discriminate the two tasks, based on voxelwise activity patterns in each DMN ROI separately (see Experimental Procedures). Classification accuracy (CA) was assessed using a leave-one-run-out procedure, and expressed as accuracy minus chance (50%). The CA for each subnetwork was then computed from the average CA of each ROI in the subnetwork.

Separately for within and between-domain task pairs, Figure 4 shows mean values of CA minus chance for each DMN subnetwork, along with results of a similar analysis for the MD network. T-tests showed classification accuracy significantly above chance for between-domain task pairs in all DMN subnetworks as well as the MD network (core: t(23) = 2.99, p < 0.01; MTL: t(23) = 3.95, p < 0.01; dmPFC: t(23) = 2.93, p < 0.01; MD: t(23) = 3.84, p < 0.01). Only in MD regions was classification accuracy of within-domain task pairs significantly above chance (t(23) = 4.42, p < 0.01). Paired t-tests revealed a significant increase in classification accuracy for between domain task pairs compared to within-domain task pairs in core and MTL subnetworks (core: t(23) = 2.59, p < 0.02; MTL: t(23) = 3.38, p < 0.01). A two-way repeated measures ANOVA was carried out with the factors network (DMN, MD) and task pair similarity (within-domain, between-domain) to test for an interaction. Data were averaged over core, MTL and dmPFC subnetworks to obtain DMN values. Significant main effects of network (F(1,23) = 8.86, p < 0.01) and condition (F(1,23) = 4.86, p < 0.05) were found, as well as a significant interaction effect (F(1,23) = 4.85, p < 0.05).

**Figure 4.**
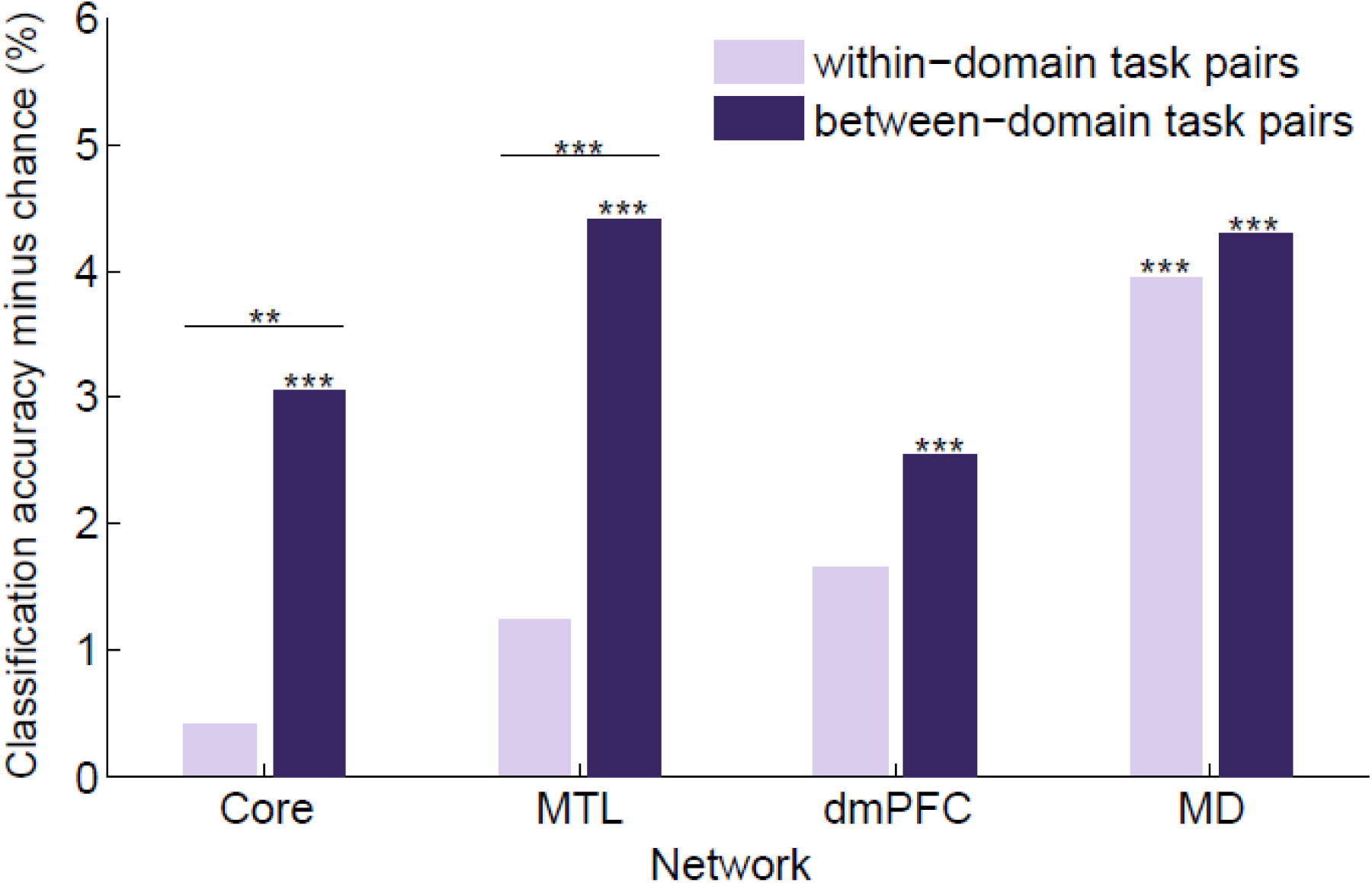
Average classification accuracies minus chance for within-domain (light bars) and between-domain (dark bars) task pairs for each DMN subnetwork and the MD network. Significant classification accuracy above chance (p < 0.05), as well as significant paired t-tests between within-domain task pairs and between-domain task pairs, are indicated with * = p < 0.05, ** = p < 0.02, *** = p < 0.01.

### Individual differences in classification accuracy predict reaction time

Finally, we related neural discrimination to task performance, testing the hypothesis that improved neural discrimination during the cue period might be associated with decreased response time (RT). Figure 5 shows the correlation between participants’ median task RT for correct trials and mean classification accuracy for each network, where classification accuracy per network was computed from the average CA of each ROI in each network. In line with the hypothesis, for MD regions there was a significant negative correlation (r = -0.55, p < 0.01), with a weaker but significant effect also seen in the DMN (r = -0.43, p < 0.05).

**Figure 5.**
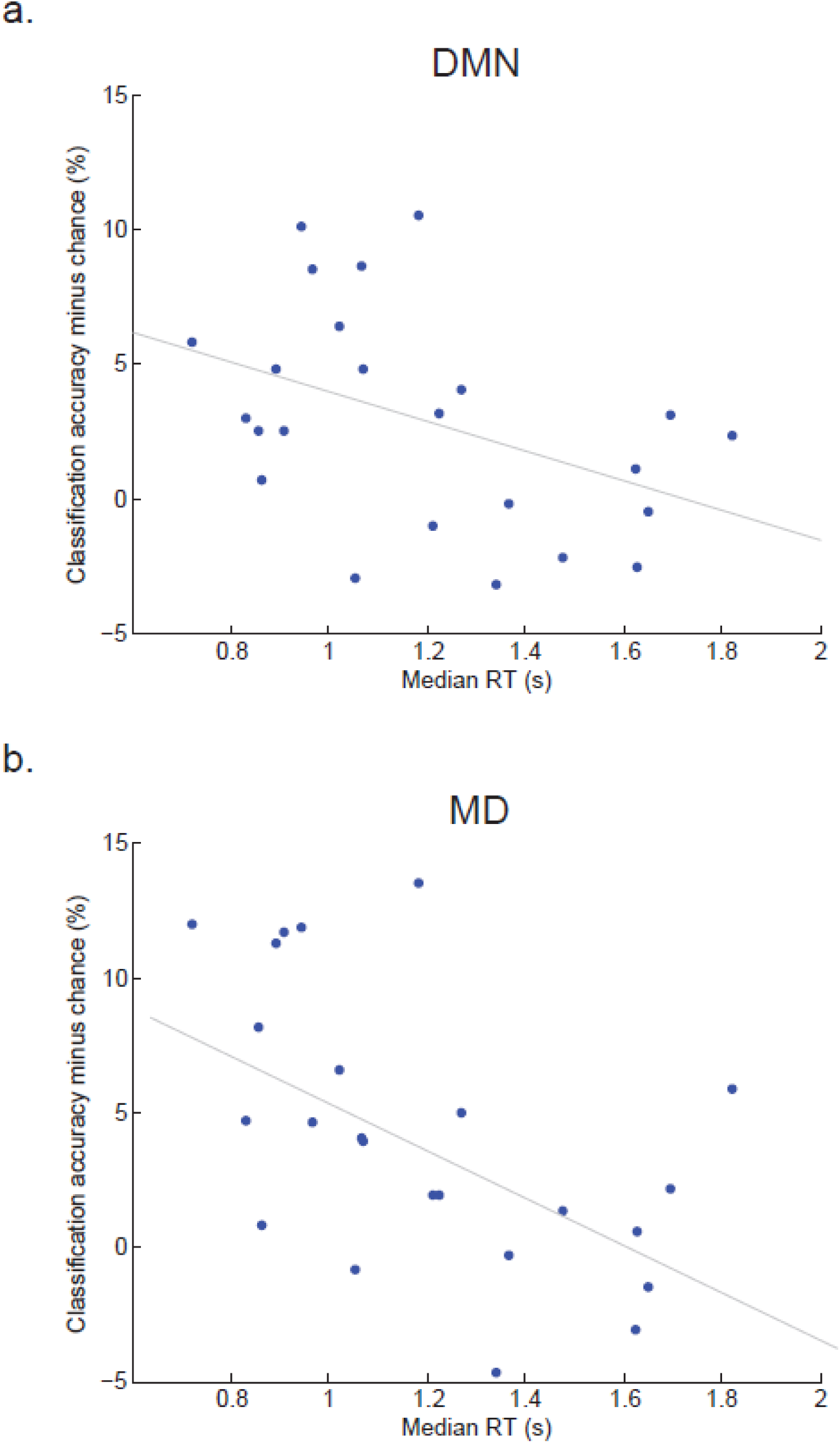
Average classification accuracy plotted against average correct reaction time (RT) for each participant. a. Mean classification accuracy over DMN ROIs. b. Mean classification accuracy over MD ROIs.

## DISCUSSION

In this study we aimed to extend a prior suggestion (Crittenden et al., 2015) that regions of the DMN can play a role in externally-directed task switching. First, we wished to replicate DMN activation during large task switches. Second, we wished to link such switch-related activity to the typical profile of DMN activation during rest. Third, our design allowed us to examine the cognitive transition from rest to task.

Our findings bring an important new perspective on DMN function. In line with Crittenden et al. (2015), we found increased activity in core and MTL subnetworks for a large change of task domain. In line with prototypical findings, we also found increased DMN activity on rest trials compared to task. Perhaps most striking, core and MTL subnetworks were also strongly active during the transition back from rest to task, with the same voxels sometimes active for both rest and restart.

In the Crittenden et al. (2015) study, it was left open whether DMN activity during a switch of task domain was caused by relaxation of the previous task set, or establishment of the next. Our introduction of rest trials indicates DMN activity for both. Activity was strong both when a previous cognitive focus was relaxed (switch from task to rest), and when a new one was established (switch from rest to the next task).

Taken together, our results are difficult to explain using theories of DMN function strictly linked to internal or self-directed cognition. Inconsistent with theories suggesting a role for the DMN in processes directed away from the current external environment (e.g. self-projection, Buckner & Carroll, 2007), we see DMN switch-related activity changes in the context of an external task. Furthermore, large DMN activity increases during short rest trials seem unlikely to be caused by immediate spontaneous mind-wandering, autobiographical memory recall or self-related cognition. Substantial activity at task restart is directly inconsistent with these theories. These results call for a reconceptualization of DMN function, addressing a role in externally as well as internally-focused cognition.

As discussed earlier, many results in the literature suggest that DMN regions represent broad features of a current scene, episode or context, including spatial surroundings (e.g. Hassabis & Maguire., 2007, 2009; Howard et al., 2014), time (e.g. Addis et al., 2007; Andrews-Hanna et al., 2010), and social aspects of self and others (Frith & Frith, 2003; D’Argembeau et al., 2005) etc. This context encoding, we suggest, can indeed be important in imagining contexts different from the current moment, but also plays a role in implementation and control of current cognition. Various possible roles might be proposed. For example, current spatial, temporal and social context could constrain what kind of behavior is currently possible, desirable or permitted. When new behavior is assembled, it makes sense that there should be reference to the options permitted by the current broad context. As another possibility, the components of current behavior must always be bound together and supported against potentially competing alternatives. Temporary association with the current, relatively stable context could be one way to achieve this.

We suggest that, as cognition unfolds, there is a constant waxing and waning of the relative prominence of context representation, and hence of DMN activity. Context representation is strong at rest, when there is little other than broad context to occupy cognition. As we progressively become embedded in the operations of a focused task, context representation may recede, perhaps because they are no longer needed. This weakening of context representations could correspond to the experience of “losing ourselves” in ongoing activity, and to well-known of anti-correlations between task-negative and task-positive regions (Fox et al., 2005a; Kelly et al., 2008). With a major switch to a new task, however, we suggest that context representation is reawakened, allowing re-reference of the new task to current surroundings. Again, this might correspond to the common experience of “becoming aware of our surroundings” when a current cognitive focus is interrupted.

Our proposal is consistent with several findings implying overlap between representations of current and internally-generated scenes. Firstly, work on the perceptuomnemonic hypothesis (Buckley et al., 2001) suggests that MTL regions are part of a perceptual processing hierarchy as well as being important for memory. DMN regions (hippocampus and parahippocampus) implicated in memory for scenes are also necessary for the perceptual processing of scenes (Lee et al., 2005; Lee et al., 2006). Studies using new multivariate techniques have found that spatial and temporal patterns of activity in further DMN regions represent scene specific contextual information when both viewing and later recalling the same episode of a TV program (Baldassano et al., 2016; Chen et al., 2017). Additionally, research using multivariate techniques suggests that information represented in DMN regions is not limited to the visuo-spatial domain. Baldassano et al. (2016) found similar patterns of DMN activity for audio-descriptions of the same television scenes, and, most relevant to our findings, research by Schuck et al. (2016) found DMN activity could decode between different non-spatial task contexts (house or face judgements). Research on grid cells further suggests that DMN regions can provide contextual structure for multiple domains of stimuli. Although initially implicated in representing the spatial structure of the environment (Doeller et al., 2010; Jacobs et al., 2013; Kraus et al., 2015), Constantinescu et al. (2016) recently found that grid cells in the entorhinal cortex and ventromedial prefrontal cortex could also represent conceptual knowledge structures, such as the neck:legs ratio in an artificial ‘bird space’.

Intriguingly, DMN regions showed some task-related patterns of activity similar to those of MD regions, with strong increases at between-domain switches, restart and rest. These results match occasional previous reports that DMN and MD activity are not necessarily anti-correlated (Christoff et al., 2009; Spreng et al., 2010; Gerlach et al., 2011; Dixon et al., 2017). In line with studies implicating MD activity in task set implementation (Dosenbach et al., 2006; Duncan, 2010, 2013; Crittenden et al., 2016), we suggest that during task restarts DMN and MD regions work together. During task restarts, DMN regions could be responsible for representation and assessment of the broad cognitive context, enabling the MD network to implement a specific task set. During large switches to a different task domain in particular, the broad task context may be briefly re-activated in order to double-check the task constraints relating to this large switch. Despite these similarities, DMN and MD networks also showed important differences. While DMN showed increased activity only for large, between-domain task switches, even small, within-domain switches recruited MD regions. Our MVPA results were consistent with this distinction, indicating only relatively coarse task representations in the DMN, while MD activity was able distinguish on a finer scale between all six tasks. In both cases, task representations were linked to performance, but the results suggest that it is MD, rather than DMN, activity that encodes and controls fine task details.

Surprisingly, we also found greater MD activity for rest compared to task cues. It is possible that the increased MD response during rest trials is due to their relative infrequency of rest cues (especially rest stays), as MD regions consistently respond to low probability inputs (Ridderinkhof et al., 2004; Frank & Cavanagh, 2014). In supplementary analyses we found greater activity in MD for rare rest stay trials compared to rest switch trials which could reflect MD region sensitivity to surprising events. Our results also have similarity to early findings on MD and DMN activity at onset and offset of task blocks (Fox et al., 2005b). Fox et al. (2005b) found transient task onset and offset activity in several MD regions (supplementary motor area, anterior cingulate, anterior insula and middle frontal gyrus) across 4 separate tasks. Interestingly, the same study also showed transient task onset and offset activity in DMN regions including the posterior cingulate, precuneus and temporo-parietal junction. Whilst DMN onset and offset activity was consistent across tasks, Fox et al. (2005b) found onset and offset activity in MD regions was more task-specific, suggesting complementary but distinct roles for DMN and MD regions in task context change. Our results also suggest that compared to DMN, MD cortex does have a somewhat different profile, showing especially high restart activity while DMN restart activity was not significantly greater than rest. This separation is consistent with the usual findings of the DMN having a relatively greater focus on rest and the MD network on task.

As task restart trials always followed rest trials, one possibility is that high DMN activity at restart reflected slow decay of neural activity, or simply a prolonged haemodynamic response, following rest. An indication that restart itself recruits strong DMN activity, over and above a sustained response to the preceding rest, comes from the data of Fox et al. (2005). In that study, transient increases in DMN activity occurred at task onsets following prolonged periods of rest. Rest activity also showed a transient component, with a peak at rest onset followed by decay as rest continued. These data, like ours, strongly link DMN activity to cognitive transitions, either from task to rest or from rest to task.

In line with the findings of Andrews-Hanna et al. (2010) and Crittenden et al. (2015), our results show substantial differences between DMN subnetworks. While activity was very similar for core and MTL subnetworks, the dmPFC network behaved quite differently, with reduced activity on switch trials, and no strong increase for rest trials. Like core and MTL subnetworks, however, the dmPFC subnetwork did show MVPA encoding of task domain. Whilst being consistent with Crittenden et al. (2015), the activity pattern in the dmPFC subnetwork is difficult to interpret. Andrews-Hanna et al. (2010) suggested that dmPFC subnetwork regions might be important for self-referential and social cognitive processes, which have been found to share considerable neural overlap (Saxe et al., 2006; Lombardo et al., 2010). One possibility is that, in a study like ours, reinstatement of social context has little involvement in cognitive transitions; potentially it is more important in everyday events, in which social context may be richer and more variable.

In summary, we have implicated the DMN in cognitive transitions, not just in internally-focused tasks but externally-focused tasks also. Just as it encodes internally-generated cognitive scenes, episodes or contexts, we suggest the DMN also encodes current, external context. Along with DMN activity, context encoding may weaken as similar cognitive operations are repeated, but reappear when major cognitive transitions call for contextual re-reference. The DMN, we suggest, is not involved simply in mind-wandering, imagination, or recollection; its contextual representations are important in shaping both internally and externally-directed cognition.

## METHODS

### Participants

28 participants (13 female), between 18-29 years old, were recruited through the Medical Research Council Cognition and Brian Sciences Unit subject panel. All participants were right handed, native English speakers, with normal or corrected to normal vision, and normal color vision. Ethics approval was granted from the Cambridge Psychology Research Ethics Committee. 4 participants (3 female) were excluded from further analysis due to technical error (3) or participant non-compliance (1).

### Task

Task events are illustrated in Figure 1. Participants were required to make same/different judgements on pairs of simultaneously presented stimuli based on a task rule. There were 3 stimulus domains with 2 task rules associated with each (male/female and old/young for face stimuli; skyscraper/cottage and inside/outside view for building stimuli; first letter and last letter for word stimuli). A further rest condition was added in which there was no task for participants to complete.

Trials of the seven tasks (including rest) were presented in a pseudorandom order. This allowed for 6 switch conditions: rest switch (rest trials preceded by task trials), rest stay (rest trials preceded by rest trials), restart (task trials preceded by rest trials), between-domain switch (task trials preceded by different-domain task trials), within-domain switch (task trials preceded by same-domain task trials) and stay trials (task trials preceded by the same task).

Each trial was split into 2 phases. In the 2 second cue phase, a colored frame was presented. Each color corresponded to a task or rest trial as represented in Figure 1. In the execution phase of task trials, two stimuli would appear and the colored frame would turn black. Participants were asked to make a “same” or “different” response, by left or right keypress, based on the task specified by the color of the frame in the cue period. There were equal numbers of “same” and “different” trials in each task and switch type. Performance was self-paced, with the imperative stimuli remaining until a key was pressed, and participants were asked to respond as quickly as possible without making mistakes. Which button corresponded to ‘same’ and ‘different’ was counterbalanced across participants. An inter-trial interval (ITI) of 1.75 seconds followed each response. To improve isolation of brain activity associated with cues – the main focus of the study – 33% of trials were catch trials, with no execution phase. Instead of an imperative stimulus, catch trials had an additional 1.2 seconds of ITI, matched to the average response time in a behavioral pilot study. This same additional ITI also followed the cue phase of rest trials.

The experiment consisted of 3 blocks of 217 trials each. Each block contained 36 stay, 36 within-domain switch, 36 between-domain switch, 36 rest switch, 12 rest stay and 24 restart trials. Of the stay, within-domain switch and between-domain switch trials, 24 contained a task execution stage and 12 were catch trials. All restart trials were full trials, including task execution. There were equal numbers of each task type for each of the task switch conditions (stay, within-domain switch, between-domain switch and restart). In addition to the above main trials, each block contained the first trial (switch type undefined), and 36 dummy trials (trials following catch trials). Dummy trials were all full trials, equally split between task types, and discarded from further analysis.

Stimuli were sourced from Wikimedia Commons and the Park Aging Mind Laboratory face database (Minear & Park, 2004). Each stimulus was positioned either side of the fixation with 3.6 degrees of visual angle from stimulus center to fixation. Each stimulus measured approximately 6.0 (width) x 4.5 (height) degrees of visual angle. The experiment was controlled using Psychophysics Toolbox for MATLAB (Brainard, 1997).

### Training

Participants were carefully pre-trained to ensure good learning of task rules. First, they were shown pairs of stimuli from each domain and asked to make same/different judgements according to each of the six task rules. Participants were then asked to learn the color of frame associated with each task rule using self-paced pen and paper memory tests. Participants were then introduced to rest trials and catch trials. To ensure fluid retrieval of task rule by frame color, a series of colored frames was presented on a monitor and participants were asked to name aloud the corresponding task rule. This portion of the training was complete when participants completed two cycles of frame colors without making a mistake. Finally, participants were given a practice block of the task. They were asked to use the cue period to prepare for the upcoming task. In the first 14 trials response feedback was given. The last 19 trials had no feedback and identical timings to the main task. Training lasted around 20 minutes, after which participants were moved into the scanner for their 3 task runs of approximately 20 minutes each. Before each run, participants were asked again to describe the rule associated with each cue color.

### Data acquisition

Images were acquired using a 3 Tesla Siemens Trim Trio magnetic resonance imaging (MRI) scanner, fitted with a 32-channel head coil. Functional magnetic resonance imaging (fMRI) acquisitions used T2*-weighted multiband Echo-Planar Imaging (multiband acquisition factor 3 for 2.5mm slices with no interslice gap, TR 1.1s, TE 30ms, flip angle 62 degrees, voxel size 2.5 × 2.5mm). T1-weighted multiecho magnetization-prepared rapid gradient-echo (MPRAGE) images were also obtained (TR 2.25s, TE 2.99ms, flip angle 9 degrees, voxel size 1 mm^3^).

### Preprocessing

Images were preprocessed using automaticanalysis (version 4) (Cusack et al., 2015) and SPM 12 (Wellcome Department of Cognitive Neurology, London, United Kingdom) for Matlab (Mathworks). The sequence of preprocessing stages involved spatial realignment of the raw EPIs, slice-time correction to the middle slice, co-registration of the functional EPI images to the structural T1-weighted image, and normalization to the Montreal Neurological Institute (MNI) template brain. For univariate analysis, functional images were then spatially smoothed using a Gaussian kernel of 10mm full-width at half-maximum. No smoothing was used for multivariate analysis.

### Regions of Interest

To stay as close as possible to the DMN subnetworks defined by Andrews-Hanna et al. (2010), we generated DMN ROIs as 8mm radius spheres around peak coordinates from that study. DMN ROIs are shown in Figure 6a. Due to the position of the bounding box, voxels covering Andrews-Hanna et al.’s (2010) original peak temporal pole coordinates were not measured; to amend this, the temporal pole volumes as used in Crittenden et al. (2015) were each dilated in volume by 2 voxels. Frontoparietal MD ROIs were taken from Fedorenko et al. (2013). MD regions (Figure 6b) included the posterior-anterior extent of the inferior frontal sulcus, frontal eye field, inferior frontal junction, anterior insula/frontal operculum, dorsal anterior cingulate/presupplementary motor area, and intraparietal sulcus. A template for these regions can be downloaded from http://imaging.mrc-cbu.cam.ac.uk/imaging/MDsystem. By using the version which separates each ROI, we were able to select only the frontoparietal ROIs.

**Figure 6.**
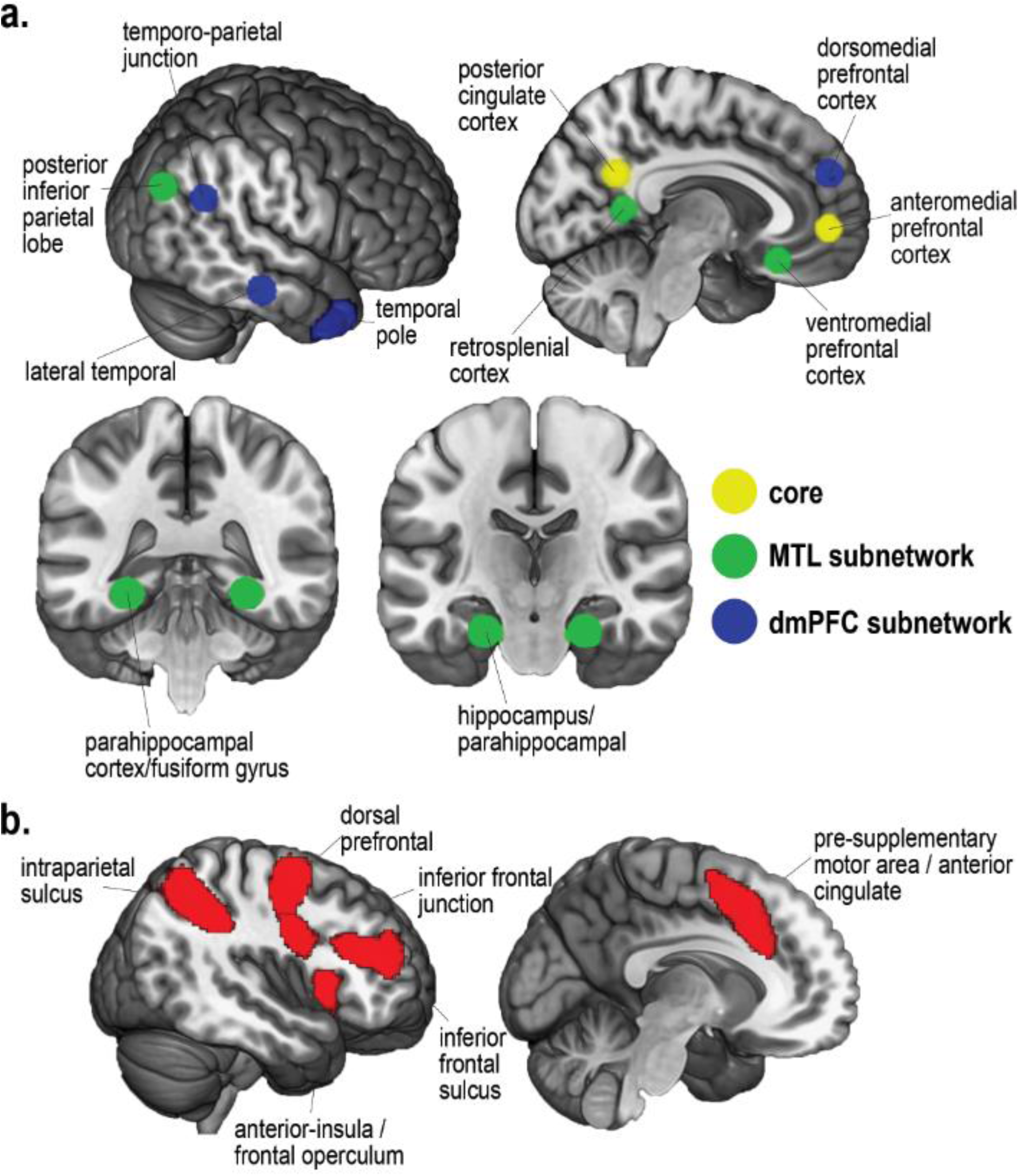
Regions of interest. a. DMN ROIs from peak coordinates presented in Andrews-Hanna et al. (2010). b. MD ROIs as presented in Fedorenko et al. (2013).

### Univariate analyses

Data for each participant were examined using the General Linear Model. Regressors were separately created for each combination of switch condition (stay, within-domain switch, between-domain switch, restart, rest switch, rest stay, dummy) by task type (gender, age, building type, viewpoint, first letter, last letter) by task phase (cue, execution). Response type (same or different) was also separated for execution phase regressors. Incorrect trials were modelled separately and discarded.

Dummy trials were also modelled and excluded from further analysis. Each regressor was modelled as delta functions convolved with the canonical hemodynamic response function, positioned at the onset of the cue periods and the middle of each execution period. Except for the restart condition, use of 33% trials with no execution phase meant that regressors could be well separated for cue and execution phases. Average contrast values were extracted for each ROI for each participant using the MarsBaR SPM toolbox (Brett et al., 2002).

### Multivariate analyses

Multivoxel pattern analysis (MVPA) was performed using the Decoding Toolbox (Christophel et al., 2012; Görgen et al., 2012). As with the univariate analysis, each regressor was modelled as a delta function convolved with the canonical hemodynamic response function, positioned at the onset of the cue period and the middle of each execution period. Incorrect trials were removed. MVPA then examined rule discrimination in patterns of cue phase activity, using the same ROIs as for univariate analysis. Prior to pattern analysis, beta values were Z-scored across tasks within each voxel of the ROI. Separate pairwise classifications were performed for each of the 15 possible task pairs (e.g. age vs. building type). Classification was carried out using a linear support vector machine (LIBSVM; Fan et al., 2005) and a leave-onerun-out approach, with the classifier trained on data from two runs and tested on the third, and results averaged over the three possible left-out runs. Classification accuracy (CA) minus chance (50%) was generated for each classification pair, for each ROI and participant. The CA for each subnetwork was then computed from the average CA across ROIs in the subnetwork.

## ACKNOWLEDGEMENTS

This work was funded by the Medical Research Council (UK) intramural program MC-A060-5PQ10.

## AUTHOR CONTRIBUTIONS

All authors contributed to project conception, experimental design, data analysis and interpretation, and writing the article. Verity Smith acquired the data.

